# Differential endothelial cell cycle status in postnatal retinal vessels revealed using a novel PIP-FUCCI reporter and zonation analysis

**DOI:** 10.1101/2024.01.04.574239

**Authors:** Ziqing Liu, Natalie T Tanke, Alexandra Neal, Tianji Yu, Tershona Branch, Jean G Cook, Victoria L Bautch

**Author notes:** Corresponding author: Victoria L Bautch, PhD, Department of Biology, CB 3280, The University of North Carolina at Chapel Hill, Chapel Hill, NC 27599 USA. current address: Department of Physiology, Medical College of Wisconsin, Milwaukee, WI 53226.

## Abstract

Cell cycle regulation is critical to blood vessel formation and function, but how the endothelial cell cycle integrates with vascular regulation is not well-understood, and available dynamic cell cycle reporters do not precisely distinguish all cell cycle stage transitions *in vivo*. Here we characterized a recently developed improved cell cycle reporter (PIP-FUCCI) that precisely delineates S phase and the S/G2 transition. Live image analysis of primary endothelial cells revealed predicted temporal changes and well-defined stage transitions. A new inducible mouse cell cycle reporter allele was selectively expressed in postnatal retinal endothelial cells upon Cre-mediated activation and predicted endothelial cell cycle status. We developed a semi-automated zonation program to define endothelial cell cycle status in spatially defined and developmentally distinct retinal areas and found predicted cell cycle stage differences in arteries, veins, and remodeled and angiogenic capillaries. Surprisingly, the predicted dearth of proliferative tip cells at the vascular front was accompanied by an unexpected enrichment for endothelial tip cells in G2, suggesting G2 stalling as a contribution to tip-cell arrest. Thus, this improved reporter precisely defines endothelial cell cycle status *in vivo* and reveals novel G2 regulation that may contribute to unique aspects of blood vessel network expansion.

## INTRODUCTION

Heterogeneity of endothelial cell responses to signaling inputs is crucial for sprouting angiogenesis and blood vessel network expansion [1–3]. For example, differential Notch signaling is important for defining tip cells vs. stalk cells in emerging sprouts [4], and Notch status also determines the response of endothelial cells to pro-angiogenic BMP signals [5]. The cell cycle is temporally regulated in actively cycling cells through G1-S-G2-M stages [3, 6, 7] and recent work shows that venous identity is linked to cell cycle gene regulation [8, 9], venous/lymphatic sprouting is regulated in G1 [10], and that cues for arterial vs venous sub-type differentiation are differentially processed in early G1 vs. late G1 [11, 12]. Cells often experience a temporary cell cycle arrest (called G0 or extended G1) and are considered quiescent [11, 13, 14], and endothelial tip cells are thought to be in a temporary cell cycle arrest due to high VEGF-A signaling [15]. Although vascular programs link to cell cycle, how cell cycle status contributes to endothelial cell heterogeneity in signaling responses and vessel network expansion is poorly understood.

The dynamic nature of cell cycle transit poses challenges for precise staging. Investigators have often taken advantage of degrons, short sequences encoding information in cis for cell cycle-mediated protein degradation [16, 17]. When linked to fluorescent reporters, time-dependent cell cycle changes are documented with spatially relevant readouts. The original FUCCI reporter distinguishes G1/G0 from S/G2/M and has been incorporated into cells and animals such as flies, fish, and mice [18, 19]. Original FUCCI mice exhibited variable reporter expression due to random transgenesis, so R26R-FUCCI2 mice were generated using mCherry-hCdt1_30-120_ and mVenus-Geminin_1-110,_ which expressed FUCCI reporters bidirectionally from the ROSA26 locus [20]. The R26-FUCCI2aR mouse ensured conditional expression of both FUCCI reporters in the same ratio [21, 22]. Newer FUCCI reporter versions utilized Cdt1_1-100_ that included the PIP degron and improved some transitions but contained other potential binding regions that may interfere with endogenous cell cycle, and reporter mice carrying these versions were not reported [23]. A more recent version utilized here has mCherry-Gem_1-110_ expression tightly confined to S and G2, along with a smaller fragment of Cdt1, Cdt_1-17_, that contains the PIP degron and whose overexpression is not predicted to affect the endogenous cell cycle; moreover, this fragment expresses precisely at the start of G2, with degradation precisely at the start of S phase [24]. This improved reporter, called PIP-FUCCI, distinguishes S, G2, and G1/G0, and it has been extensively validated in cultured transformed cells and found to reflect cell cycle status.

Here we investigated this cell cycle reporter in primary endothelial cells and in developing murine blood vessels *in vivo*. We found concordance with established cell cycle readouts and for the first time precisely define S and G2 phases in endothelial cells *in vivo*. A novel semi-automated zonation pipeline revealed that different spatial domains of the expanding retinal vascular plexus had different cell cycle stage distributions, and we documented an unexpected enrichment of endothelial tip cells in G2, suggesting how the endothelial cell cycle may integrate with vascular morphogenesis.

## MATERIALS AND METHODS

### Endothelial Cells and Imaging

HUVEC (Lonza #C2519A) were cultured according to the manufacturer’s recommendations in EBM2 (CC-3162, Lonza) supplemented with added growth factors (Bullet kit, CC-3162, Lonza, referred to as EGM2) at 37°C/5% CO_2_, and infected with PIP-FUCCI (Addgene, #118621) or H2B-CFP lentivirus at ≤ P (passage) 4. HUVEC were incubated with 1 mL of viral supernatant (prepared as described [24] in media containing 8µg/mL Polybrene (Sigma, TR-1003-G) for 4h, media was replaced for 24h, then HUVEC were seeded onto glass-bottom plates for live imaging.

Images were acquired as previously described [24]. Briefly, cells were housed in a humidified chamber (Okolabs) at 37°C/5% CO_2_ for 48h with image acquisition at 10min intervals using a Nikon Ti Eclipse inverted microscope and 20x objective lens (NA 0.75). No photobleaching or phototoxicity was observed using this protocol. Images were processed and tracked in ImageJ. Endothelial cells in the imaging field for one or more phases were chosen for phase measurements, and cells that remained in the imaging field through an entire cell cycle (mitosis to mitosis, scored by co-expression of H2B-CFP) were used for total cell cycle and movie tracks.

### Mice and breeding

All animal experiments were approved by the U. North Carolina at Chapel Hill (UNC-CH) Institutional Animal Care and Use Committee. Mice were generated and maintained on the C57BL/6J genetic background, and pups of both sexes were included in the analysis. *Cdh5-Cre^ERT2^ (Tg(Cdh5-cre/ERT2)1Rha*) mice [25] were obtained from Cancer Research UK. The new PIP-FUCCI knock-in reporter line (*C57BL/6J-Gt(ROSA)26Sor^em1(CAG-LSL-PIP-FUCCI)Vb^/Vb,* called PIP-FUCCI (PF)) was generated by the UNC-CH Animal Models Core via CRISPR/Cas9-mediated genome editing as described [26] with modifications. Briefly, the PIP-FUCCI DNA **(Fig. 1A)** was amplified via PCR, and the amplicon was cloned into a Rosa26 gene targeting construct customized for targeting with the guide RNA **(Supp. Fig 1A)**. The targeting construct contained a splice-acceptor/neomycin resistance cassette, CAG promoter, LoxP-STOP-LoxP cassette (with puromycin resistance gene), PIP-FUCCI coding sequence, Woodchuck Hepatitis Virus posttranscriptional regulatory element (WPRE), and rabbit beta-globin polyadenylation sequence. The construct, guide RNA and Cas9 protein mixture was injected into C57Bl6/J zygotes and implanted into pseudo-pregnant females. Positive founders were bred to heterozygosity for subsequent experiments. Mice carrying the PIP-FUCCI allele (*PF/PF* or *PF/+*) were bred to *Cdh-Cre^ERT2^* mice.

**Figure 1.**
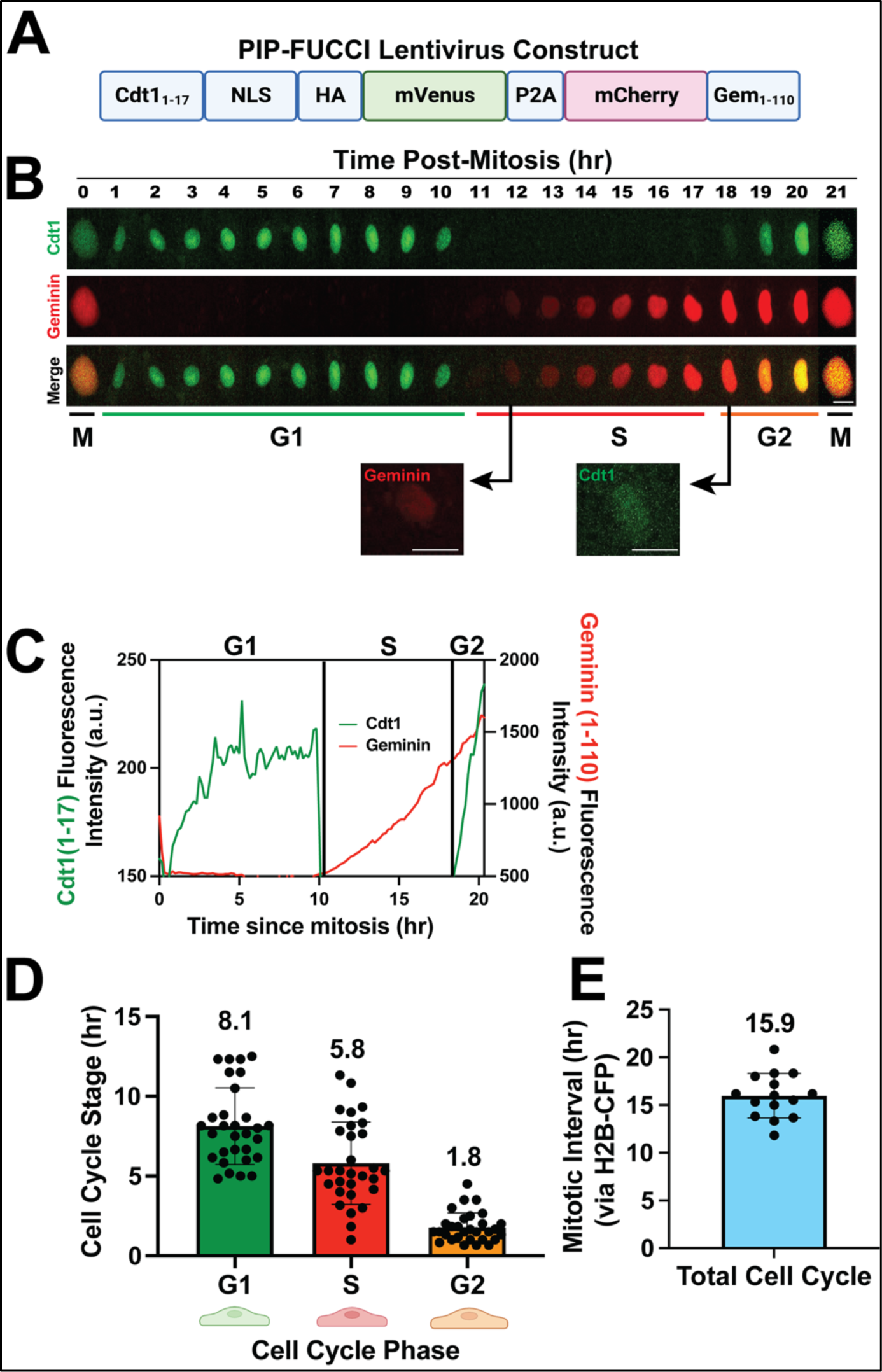
PIP-FUCCI Lentivirus reports cell cycle status in HUVEC. **(A)** PIP-FUCCI lentivirus construct. NLS, nuclear localization signal; HA, HA tag; P2A, self-cleaving peptide 2A; Gem, geminin. **(B)** Representative time-lapse images of PIP-FUCCI transduced HUVEC, showing mVenus-Cdt1_1-17_ (green) and mCherry-Geminin_1-110_ (red) expression hourly from end of cytokinesis (M) through next cytokinesis. Scale bar, 20 μm. **(C)** Quantification of PIP-FUCCI fluorescence intensity/time from cell in **B**. **(D)** Average time spent in each cell cycle state (hr). (n=30 cells per phase from 3 replicate movies) **(E).** Total endothelial cell cycle length, time between mitoses measured by H2B- CFP. (n=15 cells from 3 replicate movies).

To induce genetic deletion, 50 µl of 1mg/ml tamoxifen (Sigma T5648, 30 mg/kg body weight) dissolved in sunflower oil was injected IP into *PF/PF;Cdh5Cre^ERT2/+^*or *PF/+;Cdh5Cre^ERT2/+^* pups on P (postnatal day)1-P3 [27]. Genotyping primers amplified either wildtype *Rosa26* locus (WT-F & WT-R) or locus with PIP-FUCCI insert (PF-F & PF-R, **Supp. Fig 1B-C, Supp. Table 1**).

### Retinal staining and EdU labeling

Tamoxifen-injected pups were sacrificed at P6, eyes were collected, fixed in 4% PFA for 1h at RT (room temperature), then dissected and stored at 4°C in PBS. Staining and imaging were performed within one week of sample collection to avoid fluorescence quenching. Retinas were permeabilized in 0.5% Triton X-100 (T8787, Sigma) for 10min at RT, blocked for 1h at RT in blocking solution (5% rabbit serum (10510, ThermoFisher) and 1% BSA (A4503, Sigma) in TBST (0.1% Tween-20 in 150 mM NaCl, 50 mM Tris-HCl, pH 7.4)), then incubated with IB4 (Isolectin B4)-biotin in blocking solution overnight at 4°C. Samples were washed 3X with TBST, then incubated with Streptavidin-Alexa405 and/or ERG-Alexa647 or Ki67-Alexa647 antibody for 1h at RT (**Supp. Table 2**). Retinas were mounted with Prolong Diamond Antifade mounting medium (P36961, Life Technologies) and sealed with nail polish.

For EdU labeling, 50 µl of 3 mg/ml EdU (Thermo Fisher, A10044, 50 mg/kg body weight) dissolved in PBS was injected IP into P6 pups 2h prior to harvest [28, 29]. EdU staining was performed with the Edu-Click-iT-Alexa647 kit (C10340, Thermo Fisher) prior to staining with additional antibodies. Briefly, after the permeabilization step above, retinas were washed 2x with 3% BSA/PBS and stained with the Click-iT reaction solution at RT for 30min. Standard antibody staining for IB4 or ERG was then performed prior to mounting.

### Retina Image Analysis

Confocal images were acquired with an Olympus confocal laser scanning microscope and camera (Fluoview FV3000, IX83) using 405, 488, 561, and 640 nm lasers and 20x objective. The entire retina was imaged by collecting 5×5 or 6×6 fields and stitching. Image analysis was performed manually for EdU labeling and Ki67 staining by counting PIP-FUCCI-labeled endothelial cells from 5-8 20x images per pup. Image analysis for PIP-FUCCI/ERG co-stain followed a novel semi-automated pipeline with custom scripts in Fiji and R [30, 31].The workflow of PIP-FUCCI/ERG whole retina image analysis **(Supp. Fig 1D-E)** and vascular zonation analysis **(Supp. Fig. 2D)** was as follows. Briefly, stitched images were processed in image J to obtain the mean fluorescence intensity (MFI) of mVenus and mCherry in each endothelial cell nucleus, using ERG stain as a mask. Each nucleus was assigned a specific ID and the x y coordinates recorded. Next, vascular zones were manually drawn as polygons for each retina based on the following criteria: primary arteries (PA) and veins (PV) started from the optic disk and ended at the first bifurcation point; arterioles (Art) and venules (Ven) branched out from PA and PV and connected to a capillary bed with uniform vessel diameter; tip cells (Tip) are sprouting endothelial cells at a “tip” position on the very first row of the angiogenic front; angiogenic front capillaries (AFC) are the 3-5 rows of endothelial cells of the angiogenic front behind tip cells; mature capillaries (MC) are capillaries behind the AFC. The x-y coordinates of each vascular zone polygon were recorded. The MFI and x-y coordinates for each endothelial nucleus, and the x-y coordinates for each vascular zone were then imported into R for whole retina or zonation analysis. See supplementary information **(Supp. File 2-6)** for all custom Fiji and R scripts and for a detailed retinal zonation protocol. For zonation analysis, 2/4 retinas had no tip cells in S and 11% and 13% of tip cells scored in G2. Because indefinite numbers cannot be input for calculations and statistics, we set the G2/S ratio of these retinas in tip category at 10 (ratio was 9 and 8 for the other 2 retinas).

## RESULTS AND DISCUSSION

A new PIP-FUCCI construct that contained only the first 17 amino acids of Cdt1 linked to mVenus, along with amino acids 1-110 of Geminin linked to mCherry, was shown to precisely distinguish G1/S and S/G2 in U2OS cells [24], and here we examined primary endothelial cells expressing the reporter via lentivirus infection **(Fig. 1A)**. Live image analysis revealed sharp transitions for Cdt1_1-17_mVenus, with degradation at G1/S and re-expression at S/G2 **(Fig. 1B-C, Supp. Movie 1)**, and mitosis always followed G2 in cells imaged to this stage transition. Analysis of multiple endothelial cells revealed that G1 phase averaged 8.1h, S phase averaged 5.8h, and G2 averaged 1.8h, adding to a total endothelial cell cycle average of 15.7h **(Fig. 1D)**, in good agreement with total cell cycle transit of 15.9h, as scored by time between mitoses of H2B-CFP expressing HUVEC **(Fig. 1E)**. As described in Grant et al [24], the degradation of Cdt1_1-17_mVenus at G1/S was considered an exact measure of the start of S phase (as defined by formation of PCNA foci), and different cell lines sometimes showed a lag of Gem_1-110_ accumulation; a slight lag was documented in HUVEC, likely due to signal accumulation for Gem_1-110_. The G2/S ratio was 0.3, consistent with G2 being shorter than S in the cell cycle. Thus, primary endothelial cells regulate the PIP-FUCCI reporter in a temporal manner consistent with their cell cycle transit.

To generate a mouse carrying an inducible PIP-FUCCI allele, the PIP-FUCCI construct was placed 3’ to a standard cassette and built into the ROSA26 locus via CRISPR/Cas9-mediated insertion **(Supp. Fig 1A, see Methods)**. Mice that were either heterozygous or homozygous for the allele *C57BL/6J-Gt(ROSA)26Sor^em1(CAG-LSL-PIP-FUCCI)Vb^/Vb*, *hereafter* called PIP-FUCCI (*PF/+* or *PF/PF*) were bred to *Tg(Cdh5-cre/ERT2)1Rha* (hereafter referred to as *Cdh5-Cre^ERT2^*) mice to excise the lox-STOP-lox cassette and induce reporter expression in endothelial cells of early post-natal retinas **(Fig. 2A, Supp. Fig 1B-C)**. Overlay of the PIP-FUCCI reporter signal with Isolectin B4 (IB4, vascular-specific) or ERG (vascular endothelial-specific) staining of P6 retinal vessels showed that the PIP-FUCCI signal labeled only IB4- and ERG-positive cells, indicating endothelial cell-specific expression of the reporter in the vasculature *in vivo* **(Fig. 2B)**.

**Figure 2.**
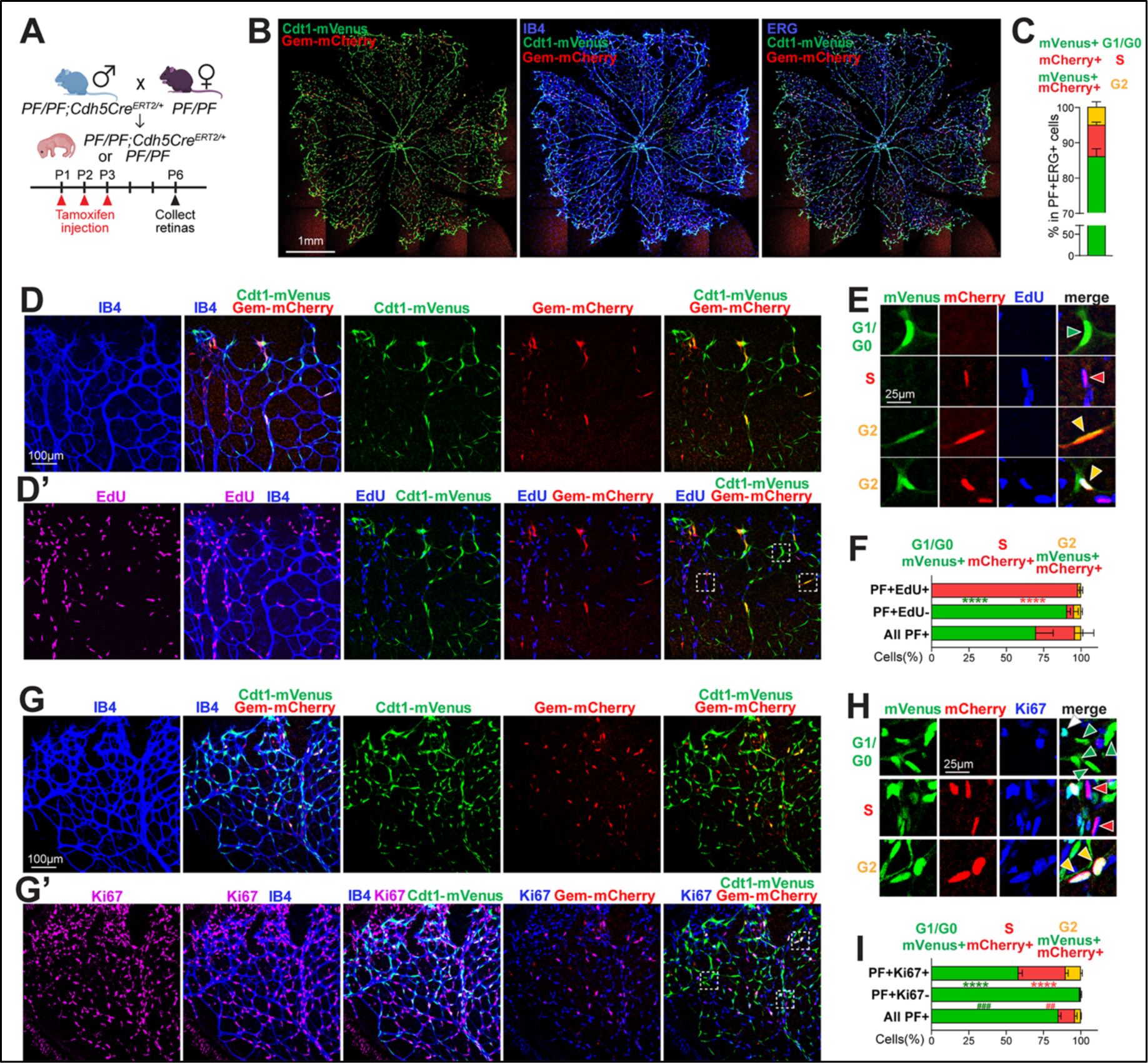
PIP-FUCCI mouse reports cell cycle status in postnatal retinal vessels *in vivo*. **(A)**. Breeding scheme and schedule for *PF/PF;Cdh5Cre^ERT2/+^* pups. **(B-C)** Representative images (scale bar, 1mm) **(B)** and quantification **(C)** of whole retina endothelial cell cycle phase analysis from PIP-FUCCI labeled P6 retinas stained for IB4 (blue) or ERG (blue). **(D-F)** Representative images **(D-E)** and quantification **(F)** of P6 PIP-FUCCI labeled retinas stained for IB4 (blue) and EdU-labeled (purple). **(E)** Representative cells (arrowheads) PIP-FUCCI labeled as in G1/G0 (green), S (red), or G2 (orange) cell cycle phase, magnified from white boxed areas in **D’ (top 3 rows of E)** and another area of the same retina **(bottom row)**. Scale bar (D) 100μm; (E) 25μm. **(F)** n = 3 pups. **** p< 0.0001 by Two-way ANOVA & Sidak’s multiple comparisons test comparing PF+EdU+ and all PF+ cells. **(G-I)** Representative images **(G-H)** and quantification **(I)** of PIP-FUCCI labeled P6 retinas stained for IB4 (blue) and Ki67 (purple). **(H)** Representative cells (arrowheads) PIP-FUCCI labeled as in G1/G0 (green), S (red), or G2 (orange) cell cycle phase, magnified from white boxed areas in **G’**. Scale bar (G) 100μm, (H) 25μm. **(I)** n = 2 pups. **** p< 0.0001 by Two-way ANOVA & Sidak’s multiple comparisons test comparing PF+Ki67+ and all PF+ cells; ### p< 0.001, ## p< 0.01 comparing PF+Ki67- and All PF+ cells.

To quantitatively analyze endothelial cell cycle in the postnatal retina with PIP-FUCCI, we created an ERG mask for all retinal endothelial cells **(Supp. Fig 1D)** that allowed for determination of mVenus and mCherry nuclear intensity in each individual endothelial cell. We then developed a semi-automated image analysis protocol (**Supp. Fig 1E**) that assigned each endothelial cell to a cell cycle phase (G1/G0, S, or G2) based on PIP-FUCCI reporter fluorescence. This analysis showed that most endothelial cells (82% for *PF/PF* homozygous retinas, **Supp. Fig 1F**) were labeled, with the small proportion of unlabeled cells likely a combination of unexcised lox-STOP-lox cassette and a few cells in very early G1 or right at the G1/S transition. Unlabeled endothelial cells were not assigned a cell cycle status or used in the quantification. Manual validation of cell cycle phase assignment using a subset of endothelial cells showed that our pipeline was accurate 96% of time (**Supp. Fig 1G**). Using this pipeline, we identified the proportion of endothelial cells in G1/G0 vs. S vs. G2 in the retina to be 86:9:5 **(Fig. 2C)**, consistent with previous reports [11]. G1/G0 is longer than S and G2, as we found in cultured HUVEC **(Fig. 1)**, although the retinal analysis revealed a lower ratio of G2 (0.06) or S (0.1) to G1/G0 cells, likely due to numerous endothelial cells that were in extended G1/G0 (quiescence) *in vivo* but not found in actively cycling cultured endothelial cells.

We next examined the relationship of the PIP-FUCCI reporter with EdU-labeling, which identifies cells in S-phase during the labeling period **(Fig. 2D-F, Supp. Fig 2A)**. Among PIP-FUCCI labeled retinal cells, mVenus+, mCherry-(green, G1/G0) endothelial cells were exclusively EdU-, showing that the Cdt1-mVenus reporter does not label S phase cells *in vivo* **(Fig. 2D-D’ Fig. 2E top row, Fig. 2F, Supp. Fig 2A)**. In contrast, mVenus-, mCherry+ cells (red, S) dominated (97.9%) PIP-FUCCI-labeled EdU-labeled cells **(Fig. 2D-D’, Fig. 2E second row; Fig. 2F)**, showing that the Gem-mCherry reporter faithfully labeled endothelial cells in S phase *in vivo*. While most (89%) of mCherry+ endothelial cells are EdU+, some mCherry+ cells score as EdU-**(Supp. Fig 2A)**, perhaps reflecting that detection of EdU labeling in tissues is likely not as sensitive as the expressed reporter. mVenus+, mCherry+ endothelial cells (orange, G2) were primarily EdU-, although a small proportion (5%) were EdU+ **(Fig. 2D’, Fig. 2E bottom row, Fig. 2F, Supp. Fig 2A)**. Since EdU labeling occurs over 2h, some endothelial cells were likely labeled in S but transitioned to G2 prior to harvest.

Ki67 reactivity identifies cells in late G1, S, and G2, although expression is heterogeneous rather than uniformly negative in G0 and G1 [32]. To further validate the PIP-FUCCI readouts, we examined the relationship of the PIP-FUCCI reporter with Ki67 **(Fig. 2G-I; Supp. Fig 2B).** Among PIP-FUCCI labeled retinal endothelial cells, mVenus-, mCherry+ (red, S) and mVenus+, mCherry+ (orange, G2) endothelial cells were almost exclusively Ki67+, consistent with our expectation that these cells were in the cell cycle *in vivo* **(Fig. 2G’, Fig. 2H second and bottom row; Fig. 2I, Supp. Fig 2B)**. In contrast, PIP-FUCCI-labeled Ki67-cells were almost exclusively mVenus+, mCherry- (green, 99.2%) **(Fig. 2G’, Fig. 2H top row green arrow heads, Fig. 2I)**, consistent with our prediction that mVenus labels G1/G0 endothelial cells. While a majority (76%) of mVenus+, mCherry-cells were Ki67- (**Supp. Fig 2B**), indicating they are likely in G0 or early G1 *in vivo*, some (24%) mVenus+, mCherry- (green) endothelial cells were Ki67+ **(Fig. 2G’, Fig. 2H top row a white arrow head, Supp. Fig 2B)**, likely due to the accumulation of Ki67 in late G1 and/or its perdurance in early G0.

Closer inspection of the retinal images revealed endothelial cells in the angiogenic front exhibited signal consistent with G1/G0 (green), S (red), or G2 (orange) cell cycle stages, while cells in the mature region were largely G1/G0 (green) **(Supp. Fig. 2C-C’’)**, suggesting spatial differences in distribution of cell cycle stages. One advantage to postnatal retinal analysis of angiogenesis is that a temporal gradient of remodeling (optic nerve outward) vs. angiogenic expansion (distal to remodeling) exists, with a clear definition of tip cells at the front vs. non-tip cells right behind the angiogenic front [4]. Spatial domains for large arteries, arterioles, and large veins and venules that form upon remodeling are also well-defined. To better understand how the cell cycle changes with time and vascular maturation, we developed a novel semi-automated zonation pipeline that identified the proportion of endothelial cells in G1/G0 vs. S vs. G2 in primary arteries (PA), Arterioles (Art), primary veins (PV), venules (Ven), mature capillaries (MC), tip cells (Tip) and angiogenic front capillaries (AFC) **(Fig. 3A, Supp. Fig 2D Supp. Files 2-6**).

**Figure 3.**
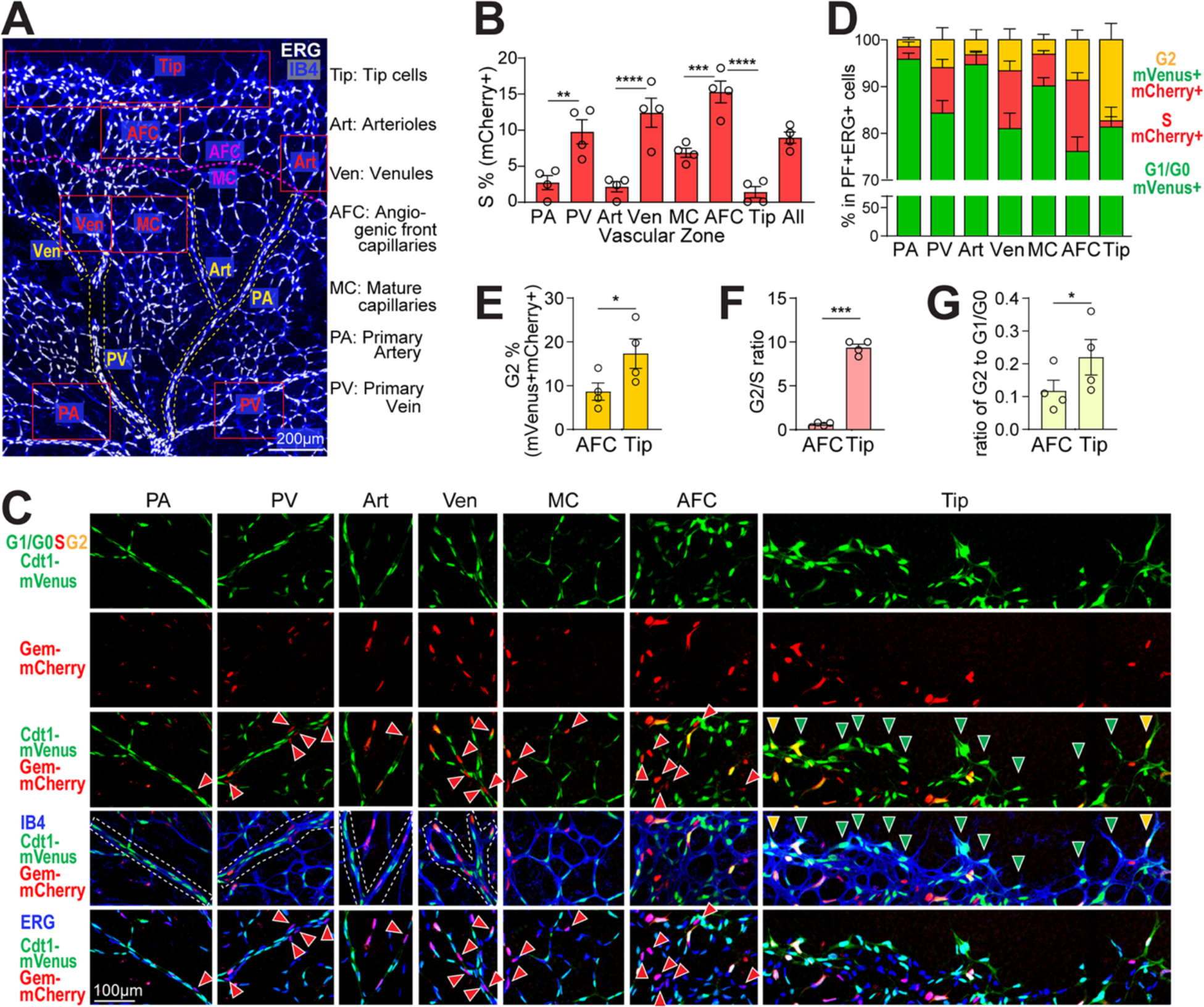
Spatial analysis reveals cell cycle differences in different retinal vascular zones. **(A)** A representative ERG (white)/IB4 (blue) labeled P6 retina image with vascular zones delineated for endothelial cell cycle analysis. Labels to right. Scale bar, 200 μm. **(B)** Quantification of % mVenus-, mCherry+ (S phase), ERG+ endothelial cells across vascular zones. **** p< 0.0001, *** p< 0.001, ** p< 0.01 by one-way ANOVA & Sidak’s multiple comparisons test. **(C-D)** Representative images (scale bar, 100μm) **(C)** and quantification **(D)** of endothelial cells (ERG+) labeled mVenus+, mCherry- (G1/G0), green arrowheads; mVenus-, mCherry+ (S), red arrowheads; and mVenus+, mCherry+ (G2M), yellow arrowheads; in retinal vascular zones. **(E-G)** Comparison of AFC and Tip cells. **(E)** Quantification of % mVenus+, mCherry+, ERG+ (G2) endothelial cells, **(F)** ratio of endothelial cells in G2 to S, **(G)** ratio of endothelial cells in G2 to G1/G0 in AFC vs. tip cells. * p< 0.05, *** p< 0.001 by paired t test. n = 4 pups for all quantifications.

As expected, the proportion of mVenus-, mCherry+ (S phase) endothelial cells was elevated in primary veins and venules compared to arterial counterparts **(Fig. 3B, C third row, PA, PV, Art & Ven**), consistent with reports that veins are more proliferative [2, 9]. Interestingly, primary arteries and arterioles consistently have a small proportion (∼2%) of S-phase labeled endothelial cells **(Fig. 3B, C third row)**, suggesting that a small proportion of arterial endothelial cells are in the cell cycle and not quiescent, consistent with a recent report showing that 2.5% arterial endothelial cells are mitotic in neonatal coronary arteries [33]. The highest S phase labeling was seen in the AFC (angiogenic front capillaries, 15%), which is significantly higher than the more mature MC zone (mature capillaries, 7%, **Fig. 3B, C third row, MC and AFC**). Very few tip cells are mVenus-, mCherry+, (S phase, 1%, **Fig. 3B, C third row Tip)**, consistent with previous report that tip cells do not usually divide [15].

Further analysis of the high-resolution zonation of cell cycle status in retinal endothelial cells revealed a surprising increase in the proportion of mVenus+, mCherry+ endothelial tip cells (orange, G2, 17.3%) compared to neighboring endothelial cells in the angiogenic front (AFC, 8.9%, **Fig. 3E)**. Additionally, while other vascular zones have 1-3 times more endothelial cells in S than G2 phase **(Fig. 3D)**, tip cells show a significantly higher G2/S ratio compared to the angiogenic front capillaries (AFC) just behind the tip (9.4 in tip cells vs. 0.6 in AFC **(Fig. 3F).** This trend is also seen in the G2/G1 ratio comparison that is significantly higher in tip cells compared to AFC (0.22 in tip cells vs. 0.12 in AFC, **Fig. 3G)**, consistent with the idea that G2-enrichment of tip cells is not simply due to decreased S phase tip cells. Thus, a portion of endothelial tip cells may be held in a G2 arrest, which suggests co-ordination of the cell cycle and cell behaviors such as sprout extension.

## CONCLUSIONS

This study validates the use of a novel PIP-FUCCI cell cycle reporter in primary endothelial cells and *in vivo*. This reporter allows for precise determination of both G1/S and S/G2 transitions not found in earlier versions, resulting from the use of the PIP degron of Cdt1_1-17_ instead of Cdt1_30-120_ in previous reporters [24]. As expected, live image analysis of primary endothelial cells revealed an average cell cycle transit time of about 16 hr, with cells on average transiting G1/G0, S, and G2 with these stages preceding mitosis. This precise cellular readout of the cell cycle will allow for a better understanding of how the cell cycle status of an endothelial cell influences its responses, as was recently shown for vascular endothelial cells in early vs. late G1 [11].

*In vivo* analysis of the PIP-FUCCI reporter took advantage of the stereotypical expansion of the postnatal mouse retinal vasculature, and we validated that the reporter readouts coincided with more traditional measures of cell cycle *in vivo*. Analysis using a novel semi-automated pipeline revealed that endothelial cells in the vascular front and veins had higher percentages of S-phase cells than arteries, and that tip cells were predominantly in either G1/G0 or G2, as predicted from their non-proliferation profile [34]. The dearth of S phase endothelial tip cells likely results from elevated levels of VEGF-A signaling [15] and is predicted to allow for migration over proliferation.

The enhanced cell cycle stage delineation of the PIP-FUCCI reporter unexpectedly led to the discovery of significant numbers of tip cells in the G2 phase of the cell cycle. This is the first reporter delineation of G2 in retinal angiogenesis, since G2-specific antibodies are rare and previous reporters did not precisely define S/G2 [18, 22]. Moreover, the ratio of G2 tip cells to tip cells in other cell cycle phases (S and G1/G0) was significantly skewed relative to nearby angiogenic front cells behind the tip cells. These relationships indicate that endothelial cells are stalled or arrested in G2 in the tip cell domain, along with numerous tip cells that are in G1/G0. The concept of a G2 stall has been described in other developmental models, such as *Drosophila* eye and sensory organ development [35, 36], as a means of co-ordinating morphogenetic movements and developmental programs. Endothelial cells in the stalk compete for the tip cell position and change positions over time [37, 38], but endothelial cells in the tip cell position rarely if ever undergo mitosis [15]. Thus, it is conceivable that an endothelial cell becomes permissive to adopt a tip cell phenotype at the S/G2 transition, and if it then moves into the tip position, a normally short G2 is extended (or stalled) to prevent mitosis during the time it resides at the tip (about 4-8 hr) [37, 38]. The prediction is that, after leaving the tip cell position, G2 cells will promptly divide, whereas G1/G0 endothelial cells will either remain quiescent or go through the cell cycle from G1 prior to mitosis.

## ACKNOWLEDGEMENTS

We thank Bautch lab members for productive discussions. We thank the UNC Animal Models Core for assistance in generating the *PIP-FUCCI* reporter mouse line. The UNC Animal Models Core Facility is supported in part by P30 CA016086 Cancer Center Core Support Grant to the UNC Lineberger Comprehensive Cancer Center. This work was supported by NIH-NHLBI R35 HL139950 and R01 GM129074 (VLB), NIH-GM083024 and GM102413 (JGC), and NIH-NHLBI 1F31HL156527 (NTT).

## SUPPLEMENTAL FIGURE LEGENDS

**Supplementary Figure 1. PIP-FUCCI reporter mice and workflow for retina analysis. (A)** Construct used to generate the PIP-FUCCI reporter mouse line. **(B)** Location of genotyping primers. **(C)** Example of genotyping results on DNA agarose gel. **(D)** Example of ERG mask. **(E)** Workflow for whole retina analysis with ERG mask. **(F)** Quantification of % PIP-FUCCI labeled cells in ERG+ endothelial cells from two *PF/PF*;*Cdh5-Cre^ERT2/+^* retinas. **(G)** Accuracy of manual vs. algorithm-assigned cell cycle phases from the same subset of endothelial cells of the same retinal images (PIP-FUCCI labeled and stained for IB4 and ERG). N = 4 pups.

**Supplementary Figure 2. Workflow for vascular zonation analysis of the retina.**

**(A)** Quantification of retinal endothelial cells EdU labeled relative to PIP-FUCCI status (related to **Fig. 2D-F)**. n = 3 pups. **** p< 0.0001 by Two-way ANOVA & Sidak’s multiple comparisons test. **(B)** Quantification Ki67+ retinal endothelial cells relative to PIP-FUCCI status (related to **Fig. 2G-I)**. n = 2 pups. **** p< 0.0001, *** p<0.001 by Two-way ANOVA & Sidak’s multiple comparisons test. **(C-C’’)** High resolution view of one representative leaflet of a *PF/PF*;*Cdh5-Cre^ERT2/+^* retina stained for IB4 and ERG. Boxed areas on far left (scale bar, 200 μm) magnified (scale bar 100 μm on middle (AFC) and far right (MC) regions. **(D)** Workflow for semi-automated vascular zonation analysis of PIP-FUCCI retinal images with ERG mask.

## SUPPLEMENTARY TABLES

**Supplementary Table 1.**
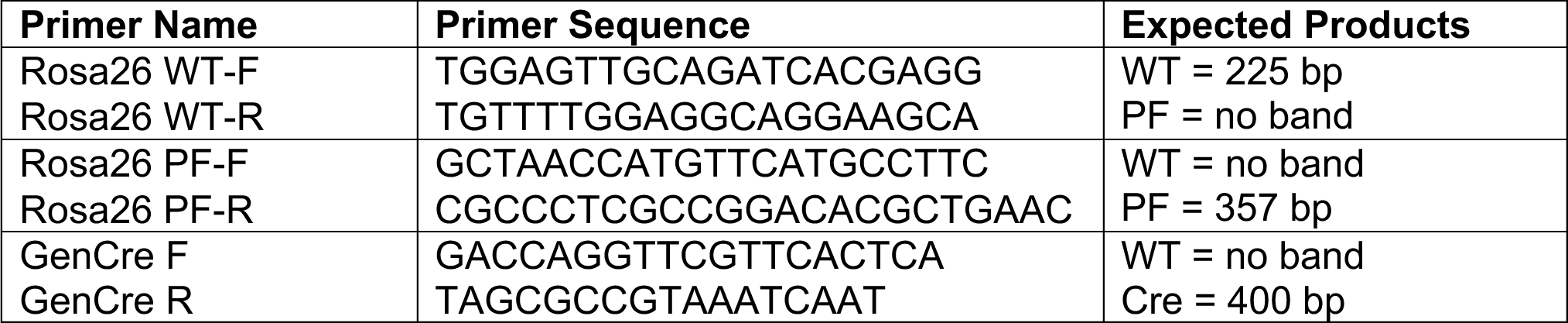
PCR Primers.

**Supplementary Table 2.**
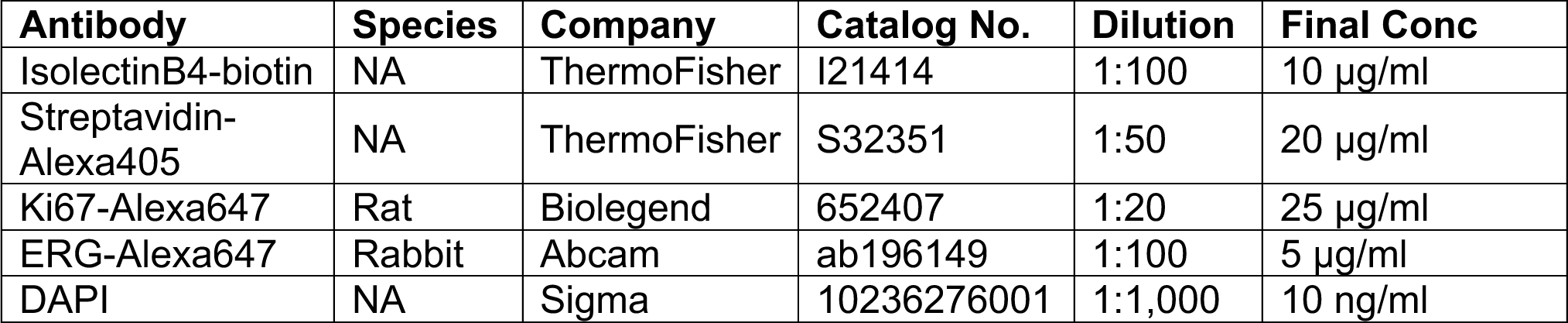
Antibodies.

## SUPPLEMENTARY ANALYSIS FILES

**Supplementary File 1.** Supplementary figures, legends, and tables.

**Supplementary File 2.** Retina_EC_PIP-FUCCI_int_extraction_by_imageJ.ijm

**Supplementary File 3.**

Retina_retrieve_xy_coordinates_from_polygon_roi_to_exclude_nonEC.ijm

**Supplementary File 4.**

Retina_retrieve_xy_coordinates_from_polygon_roi_for_vascular_zonation.ijm

**Supplementary File 5. _**R script_PIP-FUCCI_retina_EC_zonation_analysis.RMD

**Supplementary File 6.** Protocol for vascular zonation analysis of retina images.docx

## SUPPLEMENTARY MOVIE

**Supplementary Movie 1. Time-lapse imaging of representative PIP-FUCCI-expressing HUVEC.** One endothelial cell cycle with mVenus+, mCherry- (G1, green), mVenus-, mCherry+ (S, red), and mVenus+, mCherry+ (G2, orange) phases. Frames start after cytokinesis through next mitosis. Image acquisition, 1 frame/10min. Scale bar, 50 μm.

## Notes

### Competing Interest Statement

The authors have declared no competing interest.

## REFERENCES

1. Chappell, J.C., D.M. Wiley, and V.L. Bautch, Regulation of blood vessel sprouting. Seminar Cell Dev Biol, 2011. 22(9): p. 1005–1011.

2. Lee, H.-W., et al., Role of Venous Endothelial Cells in Developmental and Pathologic Angiogenesis. Circulation, 2021. 144(16): p. 1308–1322.

3. Trimm, E. and K. Red-Horse, Vascular endothelial cell development and diversity. Nature Rev Cardiol, 2023. 20(3): p. 197–210.

4. Gerhardt, H., et al., VEGF guides angiogenic sprouting utilizing endothelial tip cell filopodia. J Cell Biol, 2003. 161(6): p. 1163–1177.

5. Mouillesseaux, K.P., et al., Notch regulates BMP responsiveness and lateral branching in vessel networks via SMAD6. Nature Commun, 2016. 7(1): p. 13247.

6. Hartwell, L.H., et al., Genetic Control of the Cell Division Cycle in Yeast: V. Genetic Analysis of cdc Mutants. Genetics, 1973. 74(2): p. 267–86.

7. Minshull, J., et al., The role of cyclin synthesis, modification and destruction in the control of cell division. J Cell Sci Suppl, 1989. 12: p. 77–97.

8. Sissaoui, S., et al., Genomic Characterization of Endothelial Enhancers Reveals a Multifunctional Role for NR2F2 in Regulation of Arteriovenous Gene Expression. Circ Res, 2020. 126(7): p. 875–888.

9. Su, T., et al., Single-cell analysis of early progenitor cells that build coronary arteries. Nature, 2018. 559(7714): p. 356–362.

10. Jerafi-Vider, A., et al., VEGFC/FLT4-induced cell-cycle arrest mediates sprouting and differentiation of venous and lymphatic endothelial cells. Cell Reports, 2021. 35(11): p. 109255.

11. Chavkin, N.W., et al., Endothelial cell cycle state determines propensity for arterial-venous fate. Nature Commun, 2022. 13(1): p. 5891.

12. Fang, J. and K. Hirschi, Molecular regulation of arteriovenous endothelial cell specification [version 1; peer review: 2 approved]. F1000Research, 2019. 8(1208).

13. Coller, H.A., L. Sang, and J.M. Roberts, A New Description of Cellular Quiescence. PLOS Biology, 2006. 4(3): p. e83.

14. Johnson, M.S. and J.G. Cook, Cell cycle exits and U-turns: Quiescence as multiple reversible forms of arrest. Fac Rev, 2023. 12: p. 5.

15. Pontes-Quero, S., et al., High mitogenic stimulation arrests angiogenesis. Nature Commun, 2019. 10(1): p. 2016.

16. Alber, A.B. and D.M. Suter, Dynamics of protein synthesis and degradation through the cell cycle. Cell Cycle, 2019. 18(8): p. 784–794.

17. Coleman, K.E., et al., Sequential replication-coupled destruction at G1/S ensures genome stability. Genes & Development, 2015. 29(16): p. 1734–1746.

18. Sakaue-Sawano, A., et al., Visualizing Spatiotemporal Dynamics of Multicellular Cell-Cycle Progression. Cell, 2008. 132(3): p. 487–498.

19. Zielke, N. and B.A. Edgar, FUCCI sensors: powerful new tools for analysis of cell proliferation. WIREs Dev Biol, 2015. 4(5): p. 469–487.

20. Abe, T., et al., Visualization of cell cycle in mouse embryos with Fucci2 reporter directed by Rosa26 promoter. Development, 2013. 140(1): p. 237–246.

21. Mort, R.L., et al., Fucci2a: A bicistronic cell cycle reporter that allows Cre mediated tissue specific expression in mice. Cell Cycle, 2014. 13(17): p. 2681–2696.

22. Sakaue-Sawano, A., et al., Tracing the Silhouette of Individual Cells in S/G2/M Phases with Fluorescence. Chem & Biol, 2008. 15(12): p. 1243–1248.

23. Sakaue-Sawano, A., et al., Genetically Encoded Tools for Optical Dissection of the Mammalian Cell Cycle. Mol Cell, 2017. 68(3): p. 626–640.e5.

24. Grant, G.D., et al., Accurate delineation of cell cycle phase transitions in living cells with PIP-FUCCI. Cell Cycle, 2018. 17(21-22): p. 2496–2516.

25. Sörensen, I., R.H. Adams, and A. Gossler, DLL1-mediated Notch activation regulates endothelial identity in mouse fetal arteries. Blood, 2009. 113(22): p. 5680–5688.

26. Chu, V.T., et al., Efficient generation of Rosa26 knock-in mice using CRISPR/Cas9 in C57BL/6 zygotes. BMC Biotechnol, 2016. 16: p. 4.

27. Buglak, D.B., et al., Nuclear SUN1 stabilizes endothelial cell junctions via microtubules to regulate blood vessel formation. eLife, 2023. 12: p. e83652.

28. Honoré, T., et al., Molecular target size analyses of the NMDA-receptor complex in rat cortex. Eur J Pharmacol, 1989. 172(3): p. 239–47.

29. Zeng, C., et al., Evaluation of 5-ethynyl-2′-deoxyuridine staining as a sensitive and reliable method for studying cell proliferation in the adult nervous system. Brain Res, 2010. 1319: p. 21–32.

30. Linkert, M., et al., Metadata matters: access to image data in the real world. J Cell Biol, 2010. 189(5): p. 777–782.

31. Schindelin, J., et al., Fiji: an open-source platform for biological-image analysis. Nature Methods, 2012. 9(7): p. 676–682.

32. Miller, I., et al., Ki67 is a Graded Rather than a Binary Marker of Proliferation versus Quiescence. Cell Reports, 2018. 24(5): p. 1105–1112.e5.

33. Arolkar, G., et al., Dedifferentiation and Proliferation of Artery Endothelial Cells Drive Coronary Collateral Development in Mice. Arterioscler Thromb Vasc Biol, 2023. 43(8): p. 1455–1477.

34. del Toro, R., et al., Identification and functional analysis of endothelial tip cell– enriched genes. Blood, 2010. 116(19): p. 4025–4033.

35. Ayeni, J.O., et al., G2 phase arrest prevents bristle progenitor self-renewal and synchronizes cell division with cell fate differentiation. Development, 2016. 143(7): p. 1160–1169.

36. Meserve, J.H. and R.J. Duronio, A population of G2-arrested cells are selected as sensory organ precursors for the interommatidial bristles of the Drosophila eye. Dev Biol, 2017. 430(2): p. 374–384.

37. Arima, S., et al., Angiogenic morphogenesis driven by dynamic and heterogeneous collective endothelial cell movement. Development, 2011. 138(21): p. 4763–4776.

38. Jakobsson, L., et al., Endothelial cells dynamically compete for the tip cell position during angiogenic sprouting. Nature Cell Biol, 2010. 12(10): p. 943–953.

