## Supplemental Figures and Tables for "Differential endothelial cell cycle status in postnatal retinal vessels revealed using a novel PIP-FUCCI reporter and zonation analysis"

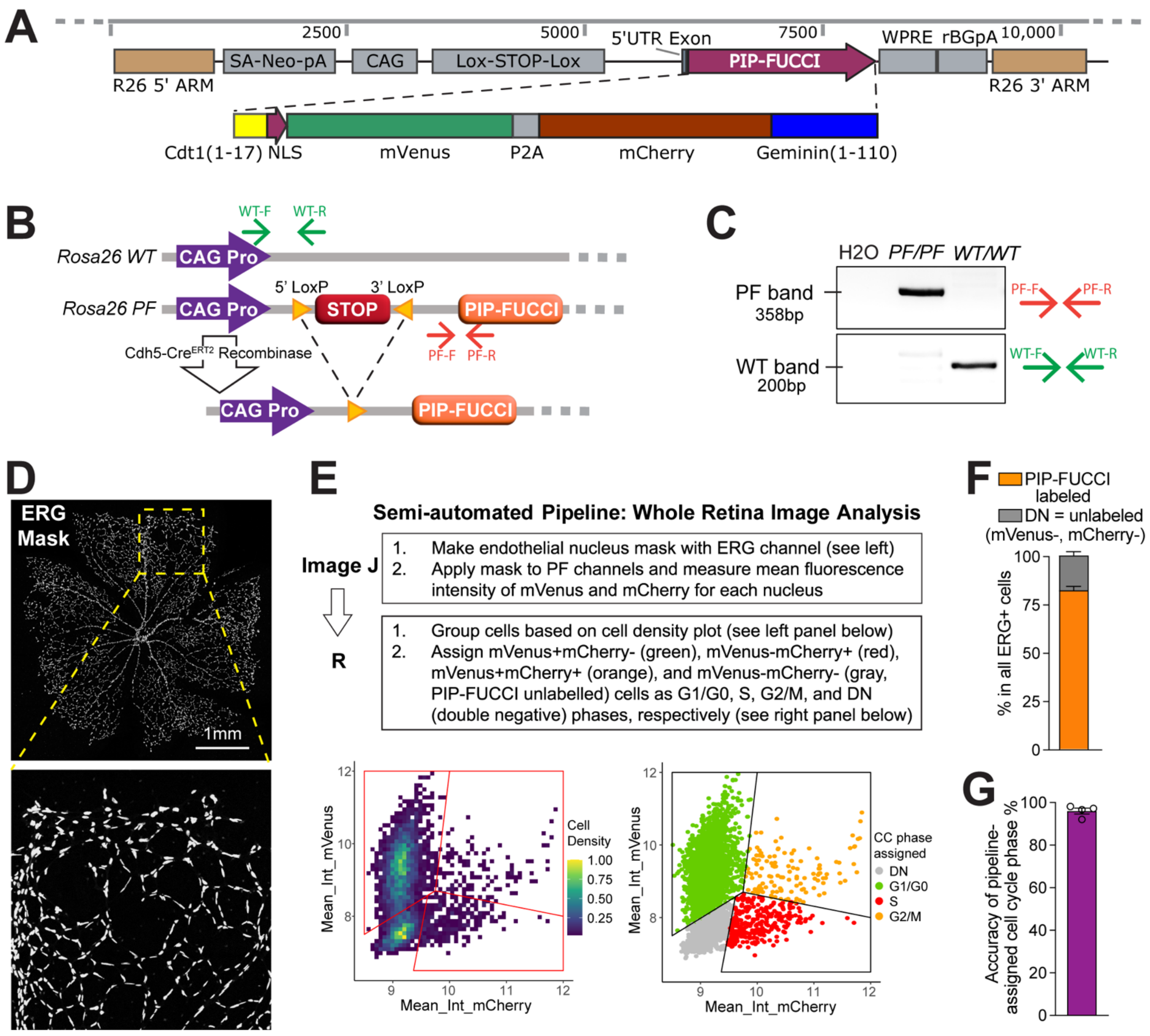


**Supplementary Figure 1. PIP-FUCCI reporter mice and workflow for retina analysis. (A)** Construct used to generate the PIP-FUCCI reporter mouse line. **(B)** Location of genotyping primers. **(C)** Example of genotyping results on DNA agarose gel. **(D)** Example of ERG mask. **(E)** Workflow for whole retina analysis with ERG mask. **(F)** Quantification of % PIP-FUCCI labeled cells in ERG+ endothelial cells from two *PF/PF*;*Cdh5-Cre*$^{ERT2/+}$ retinas. **(G)** Accuracy of manual vs. algorithm-assigned cell cycle phases from the same subset of endothelial cells of the same retinal images (PIP-FUCCI labeled and stained for IB4 and ERG). N = 4 pups.

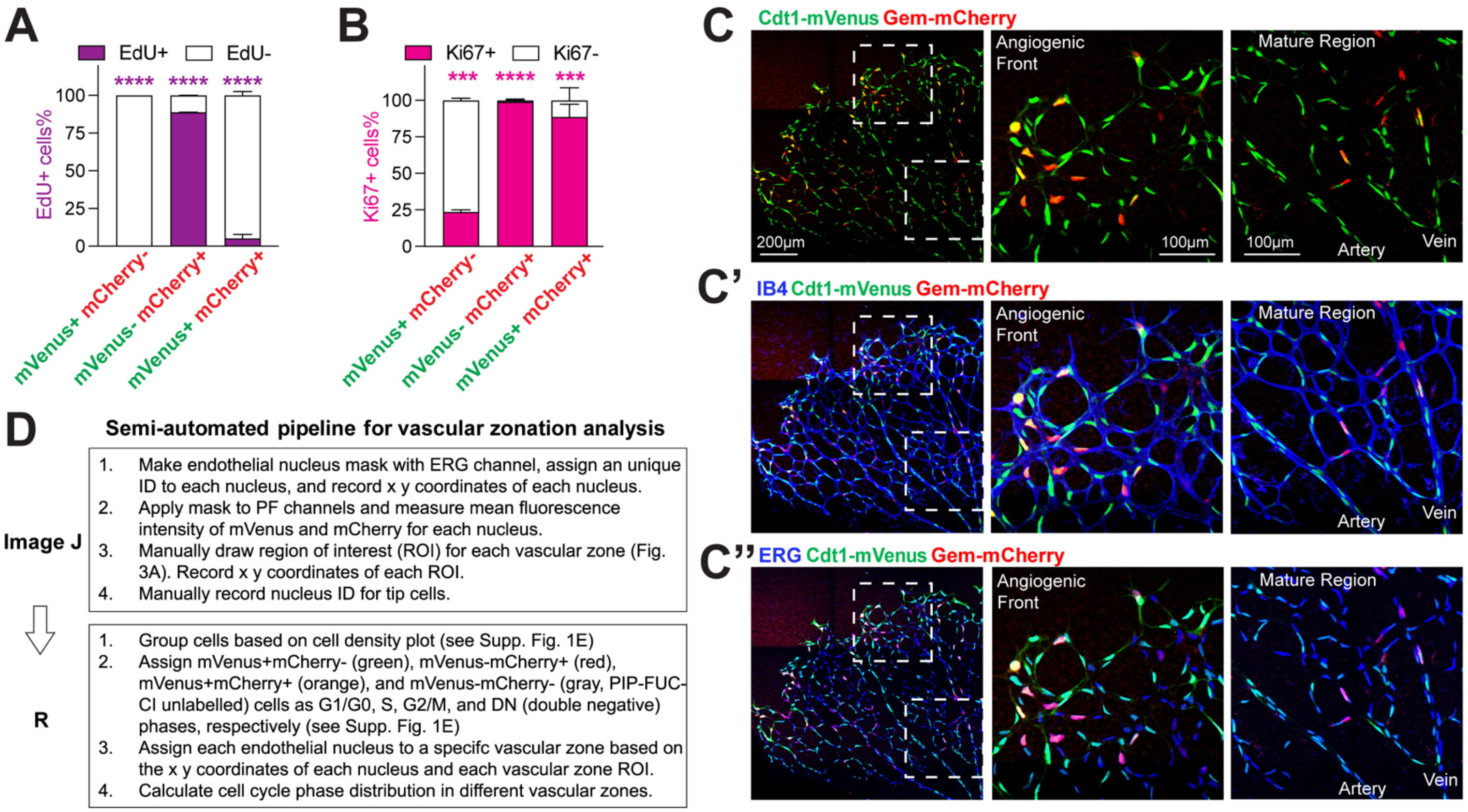


**Supplementary Figure 2. Workflow for vascular zonation analysis of the retina.**
**(A)** Quantification of retinal endothelial cells EdU labeled relative to PIP-FUCCI status (related to **Fig. 2D-F**). n = 3 pups. **** p< 0.0001 by Two-way ANOVA & Sidak's multiple comparisons test. **(B)** Quantification Ki67+ retinal endothelial cells relative to PIP-FUCCI status (related to **Fig. 2G-I**). n = 2 pups. **** p< 0.0001, *** p<0.001 by Two-way ANOVA & Sidak's multiple comparisons test. **(C-C'')** High resolution view of one representative leaflet of a *PF/PF*;*Cdh5-Cre*$^{ERT2/+}$ retina stained for IB4 and ERG. Boxed areas on far left (scale bar, 200 mm) magnified (scale bar 100 mm on middle (AFC) and far right (MC) regions. **(D)** Workflow for semi-automated vascular zonation analysis of PIP-FUCCI retinal images with ERG mask.

### SUPPLEMENTARY TABLES

#### Supplementary Table 1. PCR Primers

| Primer Name | Primer Sequence | Expected Products |
|---|---|---|
| Rosa26 WT-F<br>Rosa26 WT-R | TGGAGTTGCAGATCACGAGG<br>TGTTTTGGAGGCAGGAAGCA | WT = 225 bp<br>PF = no band |
| Rosa26 PF-F<br>Rosa26 PF-R | GCTAACCATGTTCATGCCTTC<br>CGCCCTCGCCGGACACGCTGAAC | WT = no band<br>PF = 357 bp |
| GenCre F<br>GenCre R | GACCAGGTTCGTTCACTCA<br>TAGCGCCGTAAATCAAT | WT = no band<br>Cre = 400 bp |

#### Supplementary Table 2. Antibodies

| Antibody | Species | Company | Catalog No. | Dilution | Final Conc |
|---|---|---|---|---|---|
| IsolectinB4-biotin | NA | ThermoFisher | I21414 | 1:100 | 10 μg/ml |
| Streptavidin-Alexa | NA | ThermoFisher | S32351 | 1:50 | 20 μg/ml |
| Ki67-Alexa647 | Rat | Biolegend | 652407 | 1:20 | 25 μg/ml |
| ERG-Alexa647 | Rabbit | Abcam | ab196149 | 1:100 | 5 μg/ml |
| DAPI | NA | Sigma | 10236276001 | 1:1,000 | 10 ng/ml |
